# Genomic Sequencing from Sputum for Tuberculosis Disease Diagnosis, Lineage Determination and Drug Susceptibility Prediction

**DOI:** 10.1101/2022.10.31.514503

**Authors:** Kayzad Nilgiriwala, Marie-Sylvianne Rabodoarivelo, Michael B. Hall, Grishma Patel, Ayan Mandal, Shefali Mishra, Fanantenana Randria Andrianomanana, Kate Dingle, Gillian Rodger, Sophie George, Derrick W. Crook, Sarah Hoosdally, Nerges Mistry, Niaina Rakotosamimanana, Zamin Iqbal, Simon Grandjean Lapierre, Timothy M. Walker

## Abstract

**Background:** Universal access to drug susceptibility testing for newly diagnosed tuberculosis patients is recommended. Access to culture-based diagnostics remains limited and targeted molecular assays are vulnerable to emerging resistance conferring mutations. Improved sample preparation protocols for direct-from-sputum sequencing of *Mycobacterium tuberculosis* would accelerate access to comprehensive drug susceptibility testing and molecular typing.

**Methods:** We assessed a thermo-protection buffer-based direct-from-sample *M. tuberculosis* whole-genome sequencing protocol. We prospectively processed and analyzed 60 acid-fast bacilli smear-positive sputum samples from tuberculosis patients in India and Madagascar. A diversity of semi-quantitative smear positivity level samples were included. Sequencing was performed using Illumina and MinION (monoplex and multiplex) technologies. We measured the impact of bacterial inoculum and sequencing platforms on *M. tuberculosis* genomic mean read depth, drug susceptibility prediction performance and typing accuracy.

**Results:** *M. tuberculosis* was identified from 88% (Illumina), 89% (MinION-monoplex) and 83% (MinION-multiplex) of samples for which sufficient DNA could be extracted. The fraction of *M. tuberculosis* reads from MinION sequencing was lower than from Illumina, but monoplexing grade 3+ sputum samples on MinION produced higher read depth than Illumina (*p*<0.05) and MinION multiplex (*p*<0.01). No significant difference in overall sensitivity and specificity of drug susceptibility predictions was seen across these sequencing modalities or within each sequencing technology when stratified by smear grade. Lineage typing agreement percentages between direct and culture-based sequencing were 85% (MinION-monoplex), 88% (Illumina) and 100% (MinION-multiplex)

**Conclusions:** *M. tuberculosis* direct-from-sample whole-genome sequencing remains challenging. Improved and affordable sample treatment protocols are needed prior to clinical deployment.

## Introduction

Before COVID-19, tuberculosis (TB) was responsible for more deaths than any other infectious disease. In addition, notified cases of TB dropped by 18% between 2019 and 2020, coinciding with the pandemic’s onset (1, 2). This decline is not because the caseload has reduced but because TB diagnosis capacity has been severely disrupted. Control of this much older pandemic has thereby been set back a decade. Although the impact on health systems has been an acute issue, diagnostic capacity within those systems was already under-serving population needs in many low- and middle-income settings before the pandemic.

Gold standard culture-based diagnostics remain out of reach for many patients with TB as these are dependent on centralized laboratory infrastructures, technically demanding, and expensive. Molecular platforms such as Xpert MTB/RIF Ultra™ have delivered the capacity to confirm the presence of *Mycobacterium tuberculosis* and to predict most resistance to rifampicin from primary clinical samples in settings that had previously relied on smear microscopy only (3). Nevertheless, as such targeted molecular assays are vulnerable to off-target emerging mutations and provide limited information on susceptibility to other drugs, treatment for many patients remains semi-empiric, with an increased risk of treatment failure and amplification of resistance to more drugs (4). Whole-genome sequencing (WGS) has been heralded as a potential solution for the implementation of personalized therapy, but there remains a need for a culture amplification step prior to sequencing, and solutions have so far proved stubbornly out of reach (5, 6).

Although targeted next-generation sequencing (tNGS) can identify species and lineage, predict drug susceptibility, and inform on spoligotype, it is inherently restricted in its resolution in comparative genomics for outbreak investigations compared to WGS. WGS, therefore, remains the ultimate goal (7). Here we explored how close we could get to obtaining useful diagnostic information from sequencing primary clinical samples in two high burden settings; Madagascar, a low-income country, and India, a middle-income country. We present data on samples with a range of bacillary loads, sequenced on both laboratory-confined (Illumina) and portable (Oxford Nanopore Technologies, (ONT)) sequencing platforms to assess how close we are to implementing culture-free sequencing protocols into the clinical space where they are most needed and could critically reduce TB diagnostics turnaround times.

## Methods

### Study design and sampling

The study was designed to assess direct-from-sample whole-genome sequencing on Illumina (NextSeq 500 or HiSeq 2500), and ONT MinION Mk 1B sequencing platforms, according to smear microscopy grade, with and without multiplexing on MinION flow cells. The study was conducted at the Foundation for Medical Research (Mumbai, India) and at the anti-tuberculosis dispensary Center (DAT, Antananarivo, Madagascar). The study protocol determined that the first ten patient samples should be collected from each centre for each WHO smear microscopy grade (1+, 2+, 3+). Each sample also needed to be positive for *M. tuberculosis* on Xpert MTB/RIF Ultra™ (Cepheid, USA). Six smear-negative, Xpert MTB/RIF Ultra™ negative samples were collected from each centre to use as controls. In India, these were from healthy volunteers, and in Madagascar, they were from patients with other pulmonary pathologies. All samples were divided in two, with one aliquot set up for culture and one aliquot used directly for DNA extraction on the collection day or the following day after overnight storage at 4°C. All cultured isolates were intended for WGS on Illumina. DNA extracted directly from clinical samples was divided in two for WGS on each platform. For WGS on MinION, five samples were multiplexed and five monoplexed at each centre for each smear grade, along with negative controls. In each case two, multiplexed samples were also monoplexed for cross-validation (Figure 1). Outcome measures included sequencing mean read depth, species and lineage identification, and accuracy of genotypic drug susceptibility results compared to culture-based WGS and phenotypic drug susceptibility testing results.

**Figure 1.**
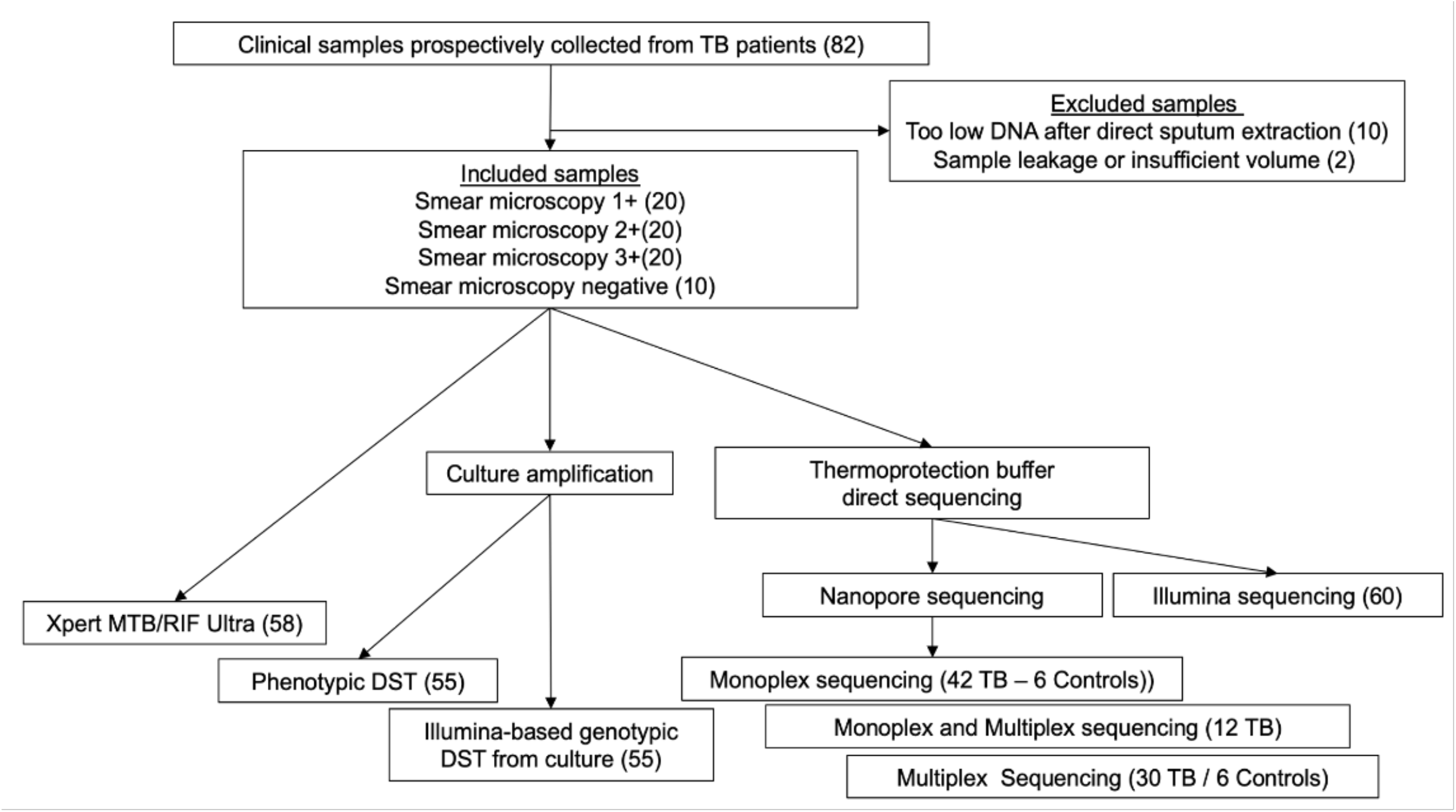
Study design including diagnostic testing, drug susceptibility testing and molecular typing assays with corresponding sample denominators.

### Sample processing

Clinical samples containing minimally 1ml of sputum were liquefied using an equal volume of thermo-protection buffer containing dithiothreitol (4 M KCl, 0.05 M HEPES buffer, pH 7.5, 0.1% dithiothreitol [DTT]) (8). 1ml aliquots were heat inactivated in 2ml screw-cap tubes at 99°C for 30 minutes. Samples were centrifuged (6,000 × g for 3 minutes) and resuspended in 100μl PBS. DNA was extracted as previously described and quantified by fluorometry (Qubit dsDNA HS assay kit, Thermo Fisher Scientific, USA) (8).

Nanopore sequencing was performed locally in each centre. DNA libraries were prepared using a ligation kit SQK-LSK 109 and sequenced on MinION devices using R9.4 flow cells. Sequencing was conducted for 48–72h, and data was acquired with MinKNOW (v19.0). In addition, the native barcoding expansion kit EXP-NBD104 (Oxford Nanopore Technologies, UK) was used for multiplexing on single flow-cells.

### Sequencing data processing

Guppy (v5.0.16) was used to de-multiplex and basecall MinION data. Illumina data were pre-processed with fastp (v0.23.2) to remove adapter sequences, trim low quality bases from read ends, and remove reads shorter than 30bp (9). Sequencing reads were decontaminated and mapped to a database of common sputum contaminants, and to the *M. tuberculosis* reference genome (H37Rv (NC_000962.3)), keeping only those reads with a mapping to H37Rv (10, 11). Decontaminated sequence files for each isolate were randomly subsampled with Rasusa (v0.6.0) to a maximum (mean) read depth of 100 (Illumina) and 150 (MinION) (12). The composition of each isolate’s sequencing data was determined by matching each read’s best mapping in the contamination alignment file to a classification category of *M. tuberculosis*, non-tuberculous mycobacteria (NTM), other bacteria, virus, human, or unmapped. Mykrobe (v.0.10.0) was used to assign lineage and predict drug susceptibility using parameters described in Hall *et. al* (10, 11, 13).

### Drug susceptibility testing (DST)

Phenotypic susceptibility testing was performed using UKMYC6 broth microdilution plate (Mumbai) and Lowenstein-Jensen (LJ) proportion method (Antananarivo) (14, 15). This was done as part of routine care and results were hence not available for all drugs. Phenotypic DST results were not used in the analysis. Culture-based WGS on Illumina results were used as reference standards as that was what we were trying to replicate by performing WGS directly from clinical samples.

### Statistics

A Wilcoxon rank-sum test was used to compare DNA concentration, read depth and accuracy of predicted outcome by smear grade and sequencing platform. Linear regression was used to assess the relationship between DNA concentration and mean read depth of the genome, having pooled the outputs of the different sequencing approaches such that one sample could have up to three results for read depth for a single DNA concentration. Fischer’s exact test was used to compare the pooled sensitivity and specificity for DST predictions to drugs of interest across each sequencing modality.

## Data availability

The decontaminated read (FASTQ) files for each isolate have been deposited in the European Nucleotide Archive (ENA) under the project accession PRJEB56100. A data sheet mapping accessions to study information can additionally be found at https://doi.org/10.6084/m9.figshare.21193588. The code used to perform all analyses in this study is available at https://github.com/iqbal-lab-org/tb-sputum-project.

## Results

Between September 2019 and January 2020, each of the two study centres collected ten sputum samples for each smear microscopy grade (1+, 2+, 3+) along with six negative controls. 58/60 samples were confirmed as *M. tuberculosis* by Xpert MTB/RIF Ultra™ and two samples from Madagascar had no result (one failed and one was not tested). 9/12 control samples were negative for *M. tuberculosis* by Xpert MTB/RIF Ultra™, and three were not tested.

All samples were set up for culture in MGIT. Twenty-three samples from Madagascar and 21 from India were positive in MGIT and could therefore undergo Illumina WGS from culture. In each case, *M. tuberculosis* and lineage were identified from the WGS data. Aliquots from the corresponding sputum samples, those that did not grow in MGIT, and negative controls were used to extract DNA directly for WGS on an Illumina platform without a culture step. Sufficient DNA was obtained for WGS from 51/60 smear-positive samples. Among these, *M. tuberculosis* was identified from 45/51 (88%) samples, with no species reported for the remaining six and a lineage from 27/51 (53%). Where a lineage or sub-lineage was called from both culture- and sputum-based WGS, 20/26 (77%) agreed at the lineage level (Table 1). All 12 negative controls were negative in culture. However, in 6/6 and 1/6 negative controls sequenced from sputum on Illumina in India and Madagascar, respectively, *M. tuberculosis* reads were identified, albeit a very small amount (<0.002% reads).

**TABLE 1.** Study samples Included number of samples, samples for which Mtb was detected and for which lineage was achieved for each combination of sequencing technology and sequencing approach (culture vs sputum). Mtb; *Mycobacterium tuberculosis*, WGS; Whole Genome Sequencing.

|  |  |  |  | Madagascar |  |  | Mumbai |  |  | Total |  |
| --- | --- | --- | --- | --- | --- | --- | --- | --- | --- | --- | --- |
|  |  |  |  | WHO smear positivity |  |  | WHO smear positivity |  |  | n | % |
|  |  |  |  | + | ++ | +++ | + | ++ | +++ |  |  |
| Illumina | culture | Samples collected |  | 10 | 10 | 10 | 10 | 10 | 10 | 60 |  |
|  |  | Culture positive and WGS result |  | 6 | 7 | 10 | 5 | 8 | 8 | 44 |  |
|  |  | Mtb detected from WGS |  | 6 | 7 | 10 | 5 | 8 | 8 | 44 | 100 |
|  |  | Lineage determined from WGS |  | 6 | 7 | 10 | 5 | 8 | 8 | 44 | 100 |
| Illumina | sputum | Samples sequenced |  | 10 | 10 | 9 | 5 | 8 | 9 | 51 |  |
|  |  | Mtb detected from WGS |  | 7 | 9 | 7 | 5 | 8 | 9 | 45 | 88 |
|  |  | Lineage called |  | 4 | 6 | 5 | 2 | 6 | 4 | 27 | 53 |
|  |  | Lineage from sputum is same as lineage from culture |  | 3/3 | 4/6 | 5/5 | 2/2 | 3/6 | 3/4 | 20/26 | 77 |
| Nanopore | sputum | monoplex | Samples sequenced | 7 | 7 | 7 | 5 | 6 | 6 | 38 |  |
|  |  |  | Mtb detected from WGS | 5 | 6 | 7 | 4 | 6 | 6 | 34 | 89 |
|  |  |  | Lineage called | 2 | 1 | 5 | 3 | 5 | 5 | 21 | 55 |
|  |  |  | Lineage from sputum is same as lineage from culture | 2/2 | 1/1 | 5/5 | 2/3 | 5/5 | 2/4 | 17/20 | 85 |
|  |  | multiplex | Samples sequenced | 5 | 5 | 5 | 0 | 4 | 5 | 24 |  |
|  |  |  | Mtb detected from WGS | 4 | 4 | 3 | - | 4 | 5 | 20 | 83 |
|  |  |  | Lineage called | 1 | 0 | 0 | - | 2 | 2 | 5 | 21 |
|  |  |  | Lineage from sputum is same as lineage from culture | 1/1 | - | - | - | 2/2 | 2/2 | 5/5 | 100 |

A subset of these primary sputum samples were further sequenced on MinION platforms. Thirty-eight were sequenced individually (one per flow cell, ‘monoplex’), and 24 were multiplexed. *M. tuberculosis* was identified from 34/38 (89%) monoplexed samples and from 20/24 (83%) multiplexed samples, and a lineage was called for 21/38 (55%) and 5/24 (21%) samples, respectively. Where a lineage or sub-lineage was called from both culture- and sputum-based WGS on MinION monoplex and multiplex, 17/20 (85%) and 5/5 (100%), respectively, agreed at the lineage level. All negative controls sequenced on MinION were negative for *M. tuberculosis*.

The reference for drug susceptibility predictions was culture-based WGS on Illumina, as that was what we were trying to replicate by performing WGS directly from clinical samples. Unfortunately, our collections included few resistant isolates, making sensitivity hard to assess. Nevertheless, resistance was correctly detected from clinical samples sequenced directly from sputum on Illumina for 6/12 (50%) isoniazid-resistant samples and 4/7 (57%) streptomycin-resistant samples. Only 1/8 (13%) rifampicin-resistant samples were detected, and 0/4 moxifloxacin-resistant samples. The number of resistant samples sequenced on MinION, either monoplex or multiplex, was lower. However, specificity was generally high, over 90% for all estimates other than for rifampicin on Illumina (83%) (Table 2; Supplementary materials 1 and 2). To assess the impact of smear microscopy grade and WGS method (Illumina, MinION monoplex/multiplex) on DST prediction from sputum, we pooled the predictions across drugs. No significant difference in overall sensitivity or specificity was seen across these sequencing modalities or within each modality when stratified by smear grade (Table 2).

**TABLE 2.**
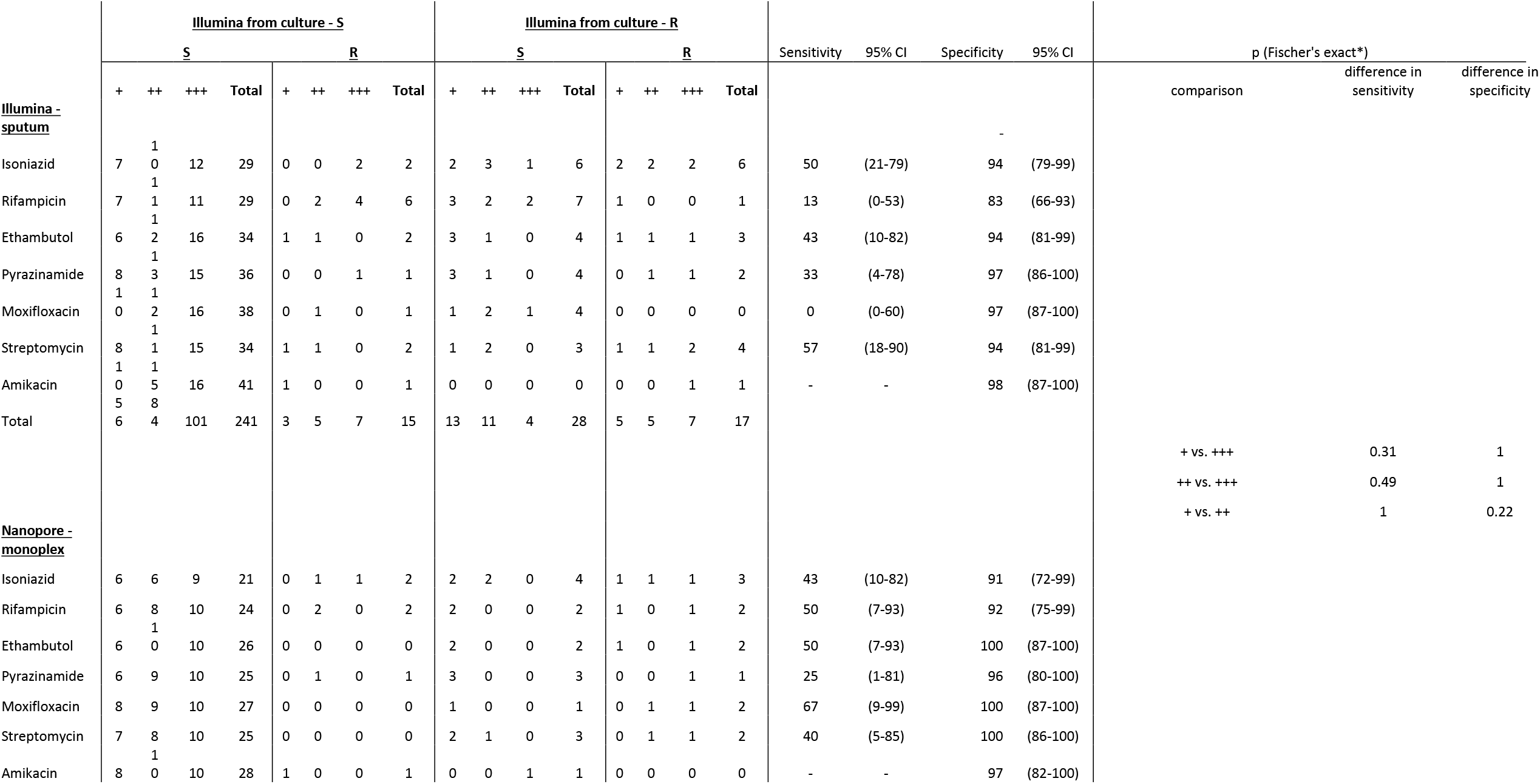
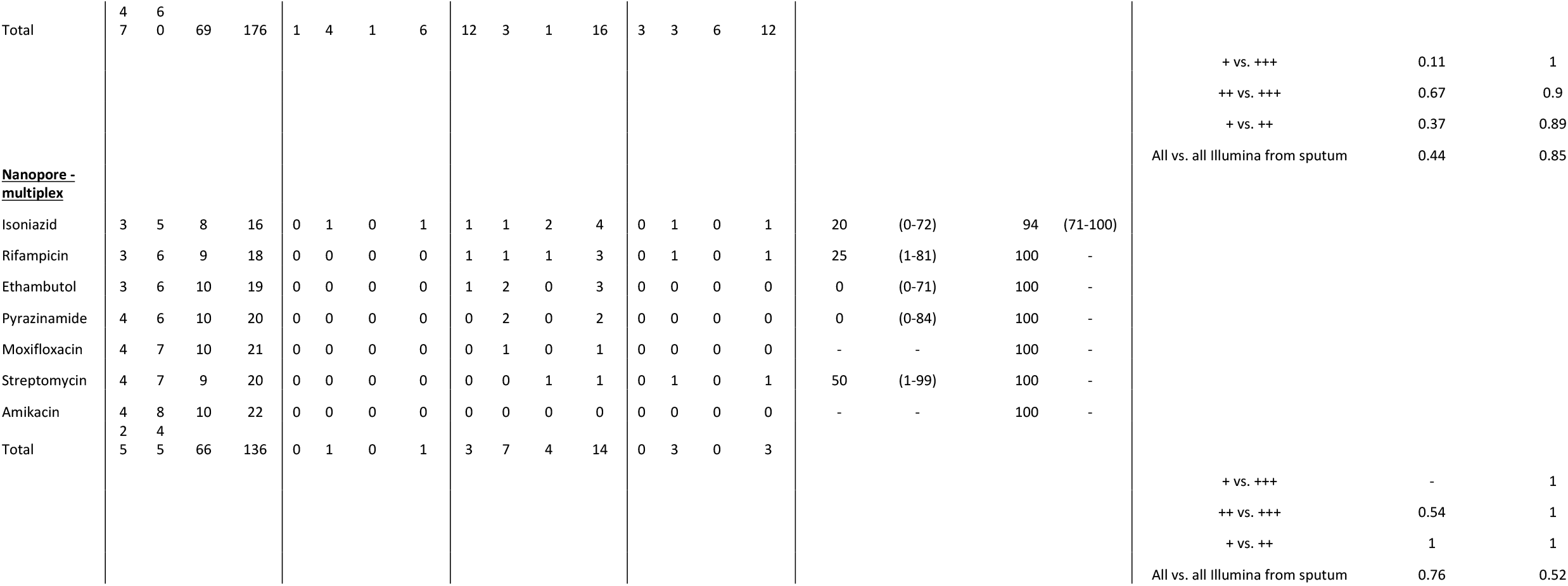
Drug susceptibility prediction agreement Drug susceptibility prediction agreement between standard Illumina sequencing from culture and each combination of sputum-based sequencing technology. R; resistant, S; susceptible* based on numbers pooled across all drugs within a given sequencing modality

Next, we sought to understand the relationship between smear grade, DNA concentration after extraction, read depth and accuracy of predicted outcomes (species, lineage and DST). There was no significant increase in DNA concentration among samples with different smear grades (Figure 2), and *M. tuberculosis* read depth did not increase with higher DNA concentrations (Figure 3). Only the smear grade 3+ monoplexed samples sequenced on MinION showed a significant increase in *M. tuberculosis* read depth - *p*<0.05 to grades 1+ and 2+ (Figure 4). Looking across sequencing modalities by smear grade, we saw that monoplexing on MinION for grade 3+ sputum samples produced the highest read depth - *p*<0.05 compared to Illumina and *p*<0.01 to MinION multiplex (Figure 4). Interestingly, the fraction of *M. tuberculosis* reads from MinION sequencing was lower than from Illumina (Figure 5), even though it had greater read depth (monoplexed).

**Figure 2.**
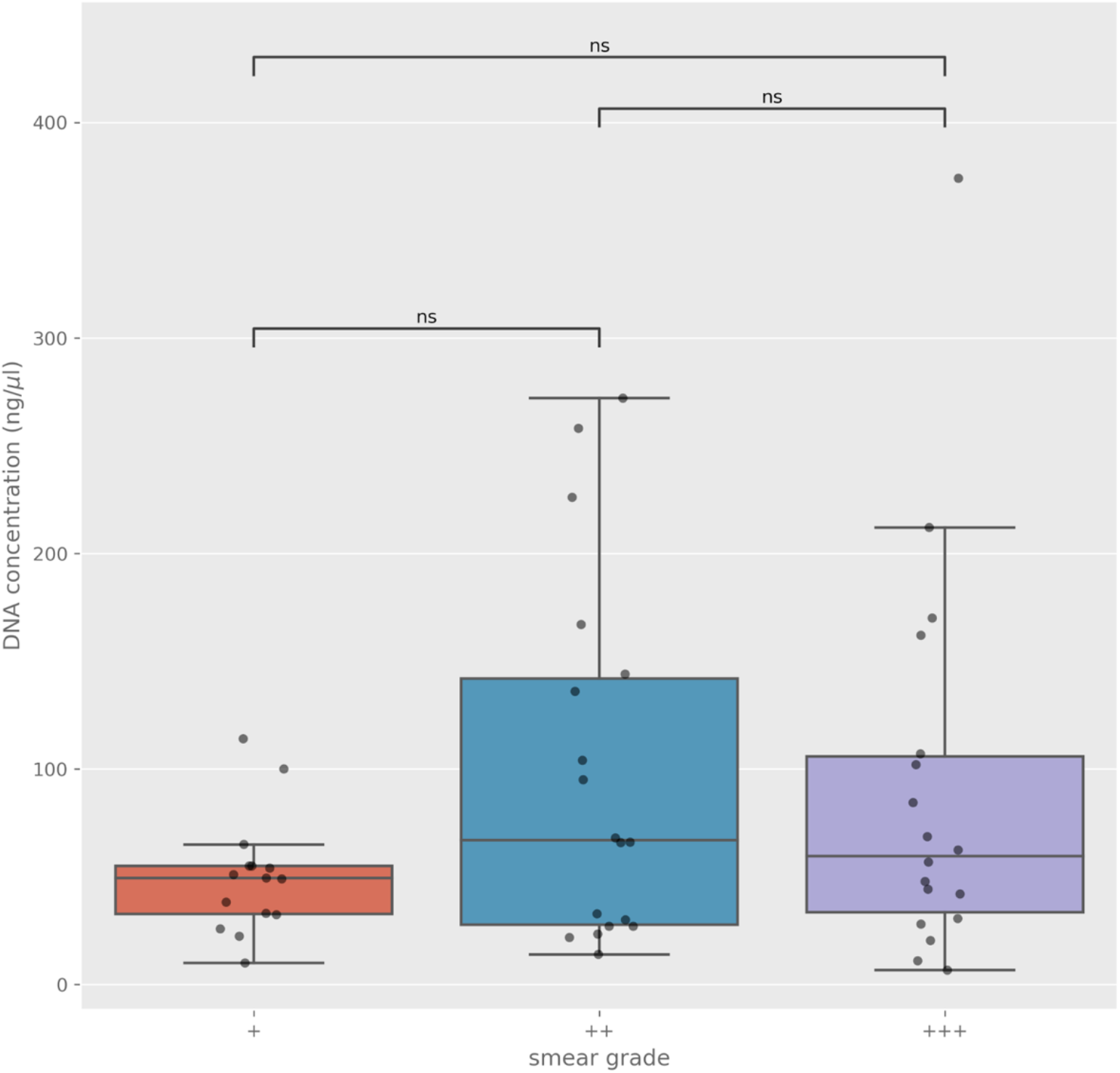
DNA concentration (y-axis) for each smear grade (x-axis). Each point represents a single sample. Annotated *p*-values were calculated with a Wilcoxon rank-sum test. ns - not significant.

**Figure 3.**
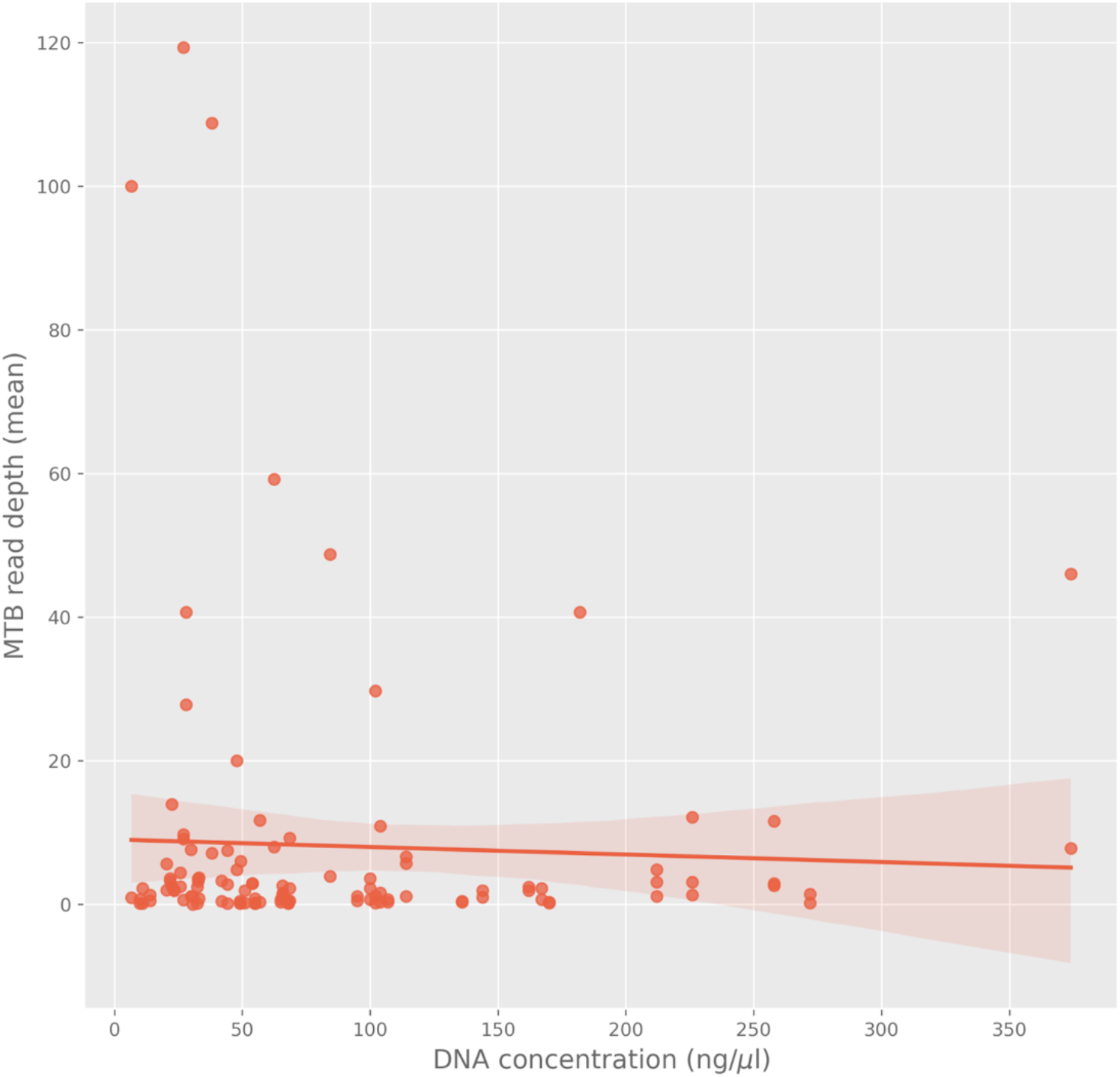
Relationship between mean read depth on the *M. tuberculosis* genome (y-axis) and the concentration of DNA extracted from the isolate (x-axis). The shaded area represents the 95% confidence interval. Each point represents a sample-sequencing strategy pair, so some samples will appear multiple times.

**Figure 4.**
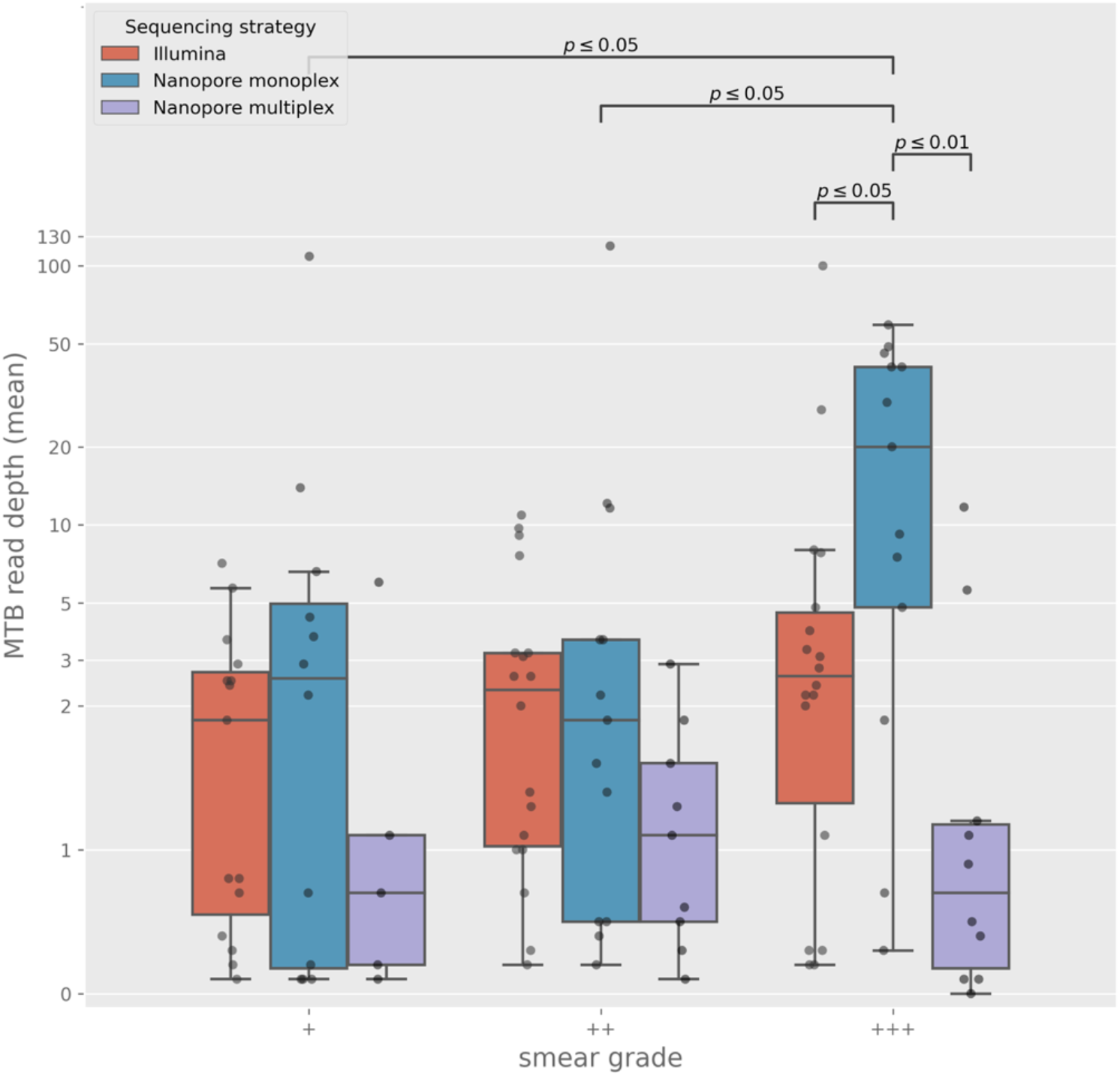
Mean read depth on the *M. tuberculosis* genome (y-axis), stratified by smear grade (x-axis) and sequencing strategy (colours). Each point represents a single sample. Annotated *p*-values were calculated with a Wilcoxon rank-sum test.

**Figure 5.**
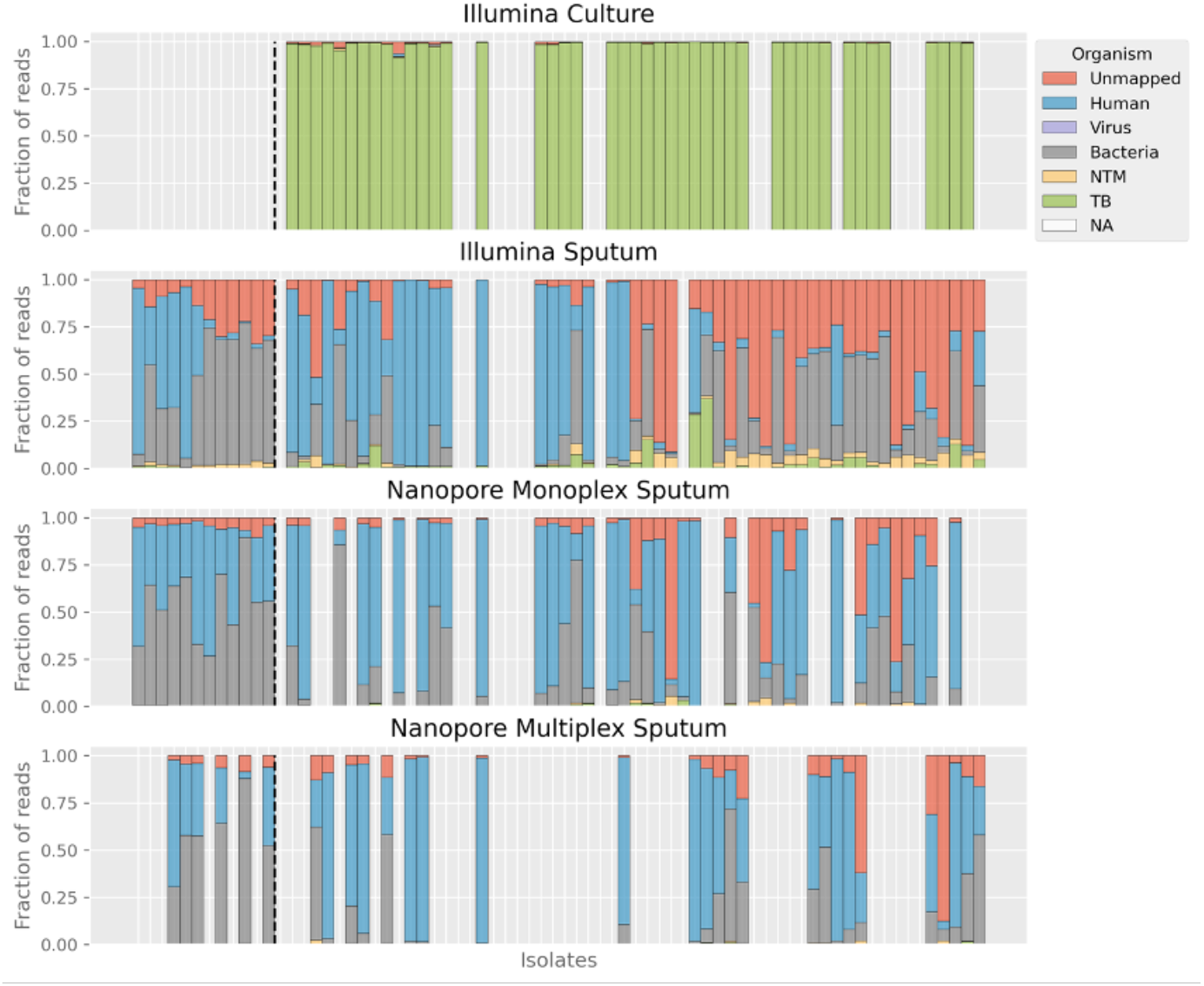
Fraction of reads (y-axis) from sequencing strategies that align to given organisms (colours). Each bar represents a single sample. Bars are vertically aligned such that the same isolate has the same place on the x-axis across the sequencing strategies. Where a bar is missing, that isolate was not sequenced with that strategy. Bars to the left of the vertical black dashed line are negative controls. Note, most sputum isolates have TB reads, but the fraction is so low it is sometimes not possible to see it. NTM - non-tuberculous mycobateria. TB - *Mycobacterium tuberculosis*. NA - sequencing not available.

Despite the limited evidence for increased *M. tuberculosis* read depth with DNA concentration or smear grade, there was clear evidence that increased read depth led to improved predictions of both species and lineage (Figure 6). For DST this was only apparent for smear grade 1+ sputum samples (*p*<0.01) but not for grades 2+ and 3+ samples (Figure 6).

**Figure 6.**
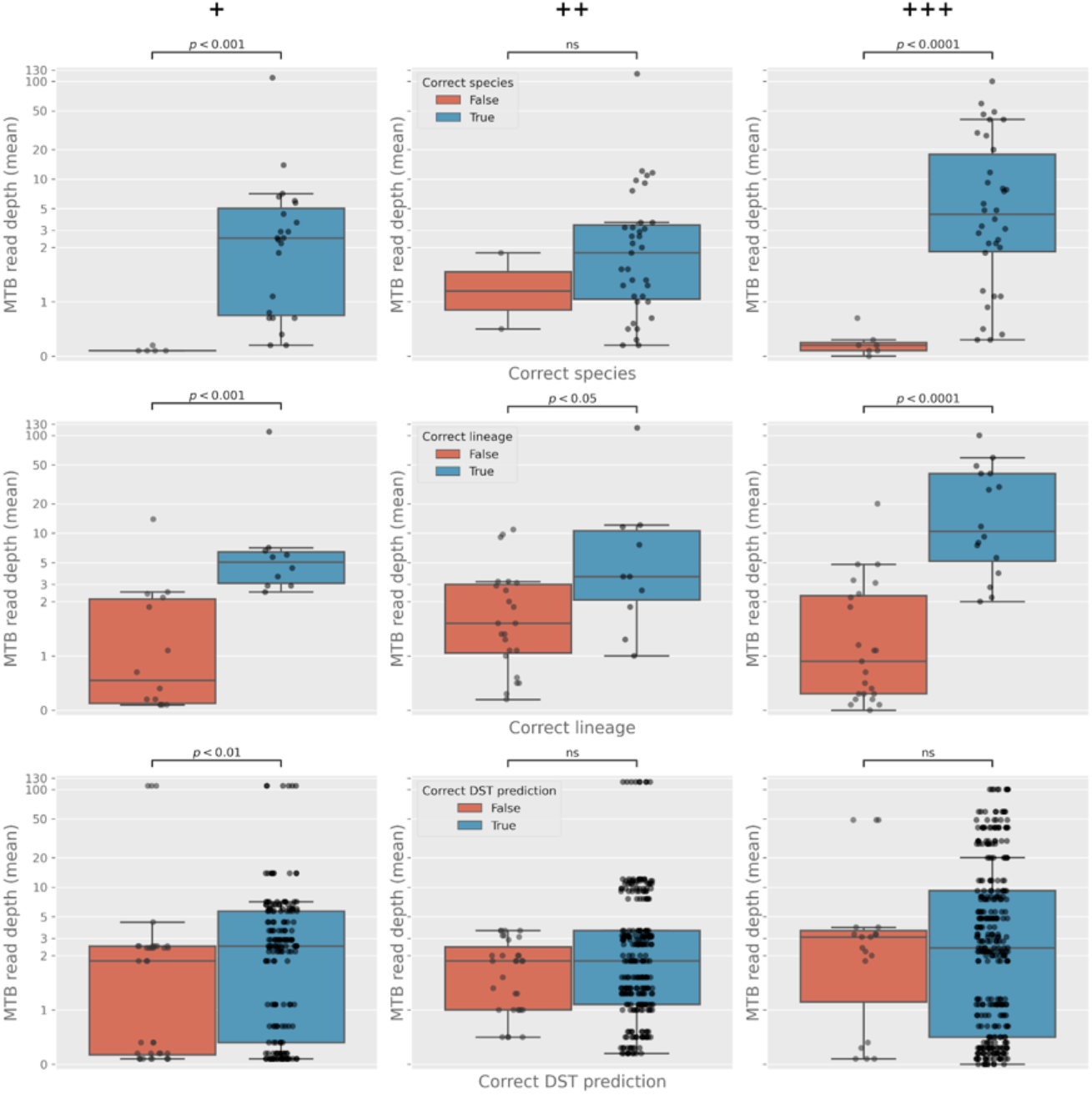
Impact of mean read depth (x-axis) on predicting species (top panel), lineage (middle panel), and drug susceptibility (DST; bottom panel). Samples are stratified by whether the prediction was correct (blue) or not (red). Note, for DST, each point is a sample-drug pair, hence why it has more points than species and lineage. Annotated *p*-values were calculated with a Wilcoxon rank-sum test.

## Discussion

This study aimed to derive clinically useful information from WGS performed on primary clinical samples in two high TB-burden settings. We compared results from Illumina and ONT MinION platforms, using monoplex and multiplex approaches on MinION, across various sputum smear microscopy grades. Using Illumina WGS data from the cultured isolates as a reference, we assessed how well we could replicate culture-based WGS predictions for *M. tuberculosis* species, lineage, and drug susceptibility from sputum samples.

Given how challenging direct-from-sample WGS still is for *M. tuberculosis*, it is unsurprising that the results are not yet good enough for clinical deployment. Although species could be correctly called for 83-89% of samples, a lineage could only be called for 21-55%, although it was mostly called correctly in those samples. Drug susceptibility predictions were also challenging, with sensitivity often around 50% or lower and no drugs performed consistently well. Specificity was generally high - only Illumina sputum rifampicin predictions (83%) had a value lower than 90% - but that is likely a function of the predominance of susceptible strains in the collection. We aimed to assess an inexpensive approach which could be deployed in low- and middle-income settings. Brown et al. previously used a biotinylated RNA baits approach and recovered over 98% of the *M. tuberculosis* genome from 20 of 24 (83%) of smear-positive, culture-positive sputa included in their study (16). This approach outperformed ours but is significantly more expensive and hence not foreseen as being widely implemented in the context of high TB.

It is a reasonable assumption that the per-sample cost of WGS will likely be affected by whether the samples are monoplexed or multiplexed on a MinION. We unsurprisingly obtained greater read depth by monoplexing than multiplexing. However, we did not find that read depth from multiplexed MinION samples differed from those sequenced on Illumina, suggesting that with improved approaches to enriching mycobacterial DNA, there may be a future for multiplex WGS on MinION. The number of negative controls from which *M. tuberculosis* reads were identified after Illumina sequencing was concerning, especially if culture-free sequencing would be used simultaneously for TB diagnosis and DST. It is clear that the risk of cross-contamination is substantial and will only grow as higher concentrations of DNA are used.

Interestingly, there was minimal evidence of a relationship between smear grade, DNA concentration when extracting directly from sputum samples, and mean read depth across the genome. Read depth was nevertheless a decisive factor in determining the correct species, lineage, and to some extent, drug susceptibility predictions. The implication is that it might be challenging to use the smear grade or DNA concentration after extraction to predict whether valuable results will likely emerge from sequencing a given specimen.

One of the main challenges with performing WGS directly from clinical samples is that the amount of extractable mycobacterial DNA is tiny compared with that from human and non-mycobacterial microorganisms. This was reflected in the meagre fraction of reads mapping to *M. tuberculosis*. Culture amplifies this fraction enormously with the cost of slowing the diagnostic process. Targeted next-generation sequencing is an alternative approach to obtaining species, lineage and DST predictions by amplifying multiple molecular targets (17). Although it can also be used for spoligotyping, the resolution of spoligotyping for comparative genomics is intrinsically limited compared to that obtained by WGS (18, 19).

Performing WGS directly from clinical samples remains desirable despite the capabilities of tNGS. Obtaining all clinical information from a single assay in a timely, reliable and cost-effective manner would be a significant advance. We have assessed how well WGS direct-from-sputum performs in our hands when applied in high-burden, low- and middle-income settings. We focussed on predicting species, lineage and DST rather than performing comparative genomics and show that even for these relatively more straightforward tasks, there remains much room for improvement. Whole genome amplification approaches may yield better results than reported here and should sensibly be pursued in the future.

## Acknowledgement

MBH was supported, in part, by an Australian Government Medical Research Future Fund (MRFF) grant (2020/MRF1200856). SGL is supported by a Junior 1 Salary Award from the Fonds de Recheche Santé Québec.

