## Supplementary Figure 1 - 2 for "Genomic Sequencing from Sputum for Tuberculosis Disease Diagnosis, Lineage Determination and Drug Susceptibility Prediction"

### Supplementary Materials

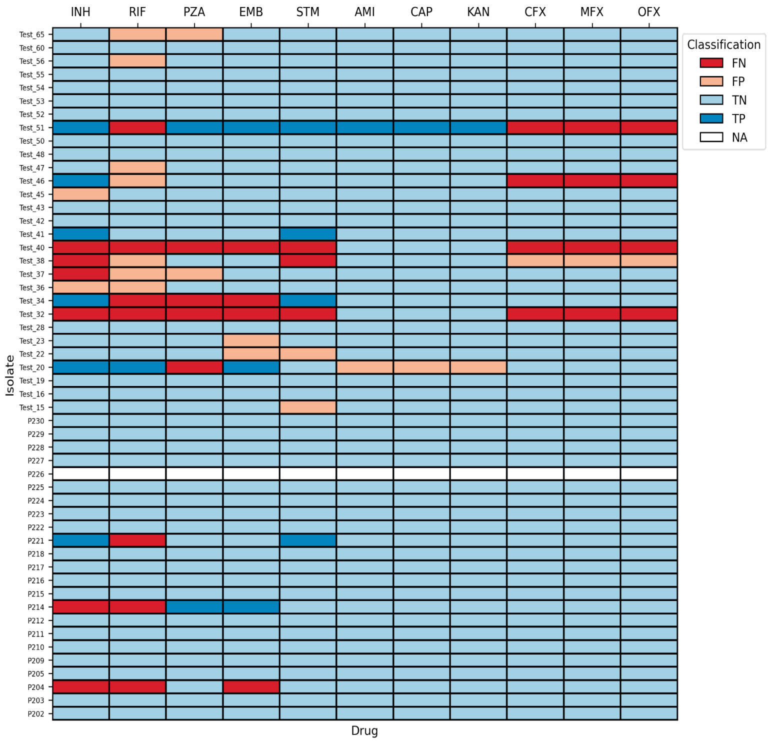


Supplementary material 1 - Drug susceptibility testing accuracy of Illumina-based direct-from-sputum sequencing using culture-based WGS on Illumina results as the reference standard. FN - false negative. FP - false positive. TP - true positive. TN - true negative. NA - data not available.

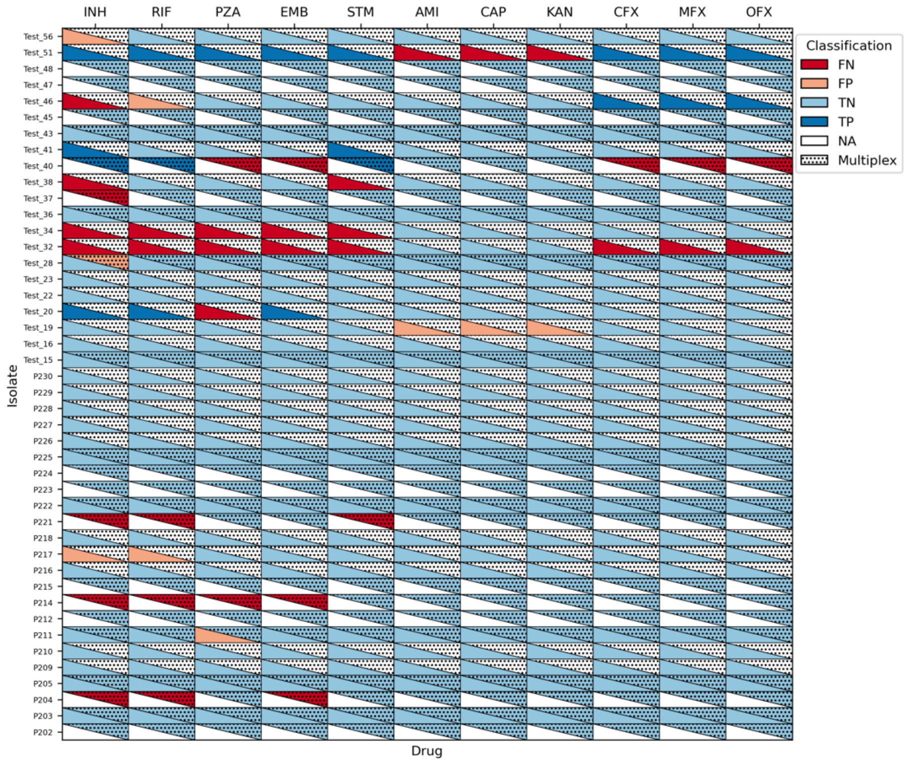


Supplementary material 2 - Drug susceptibility testing accuracy of nanopore MinION-based direct-from-sputum sequencing using culture-based WGS on Illumina results as the reference standard. For each sample, predictions from monoplexing and multiplexing (triangles with dots) are presented when available. FN - false negative. FP - false positive. TP - true positive. TN - true negative. NA - data not available.
